# MiR-384 Regulates Reelin by Inhibiting ADAMTS4 in Neuronal Cell Lines

**DOI:** 10.1101/2021.06.04.447133

**Authors:** Qiushi Cao, Bisheng Huang, Ping Wang, Gang Zhao, Min Zhao, Xiao-Ming Yu

**Affiliations:** Psychology Department, Hubei Provincial Hospital of Traditional Chinese Medicine, Wuhan 430061, China; College of Basic Medicine, Hubei University of Chinese Medicine, Wuhan 430065, China

**Keywords:** Reelin, miR-384, ADAMTS4, Alzheimer’s disease, neuronal cell line

## Abstract

MicroRNAs (miRNAs) are important regulators of gene expression at the post-transcriptional level. The present study aims to investigate the role of miR-384 in Reelin by regulating ADAMTS4 in neuronal cell lines. Brain tissues from Aβ_1-42_ induced mouse model of Alzheimer’s disease and the control group were collected. RT-PCR, Western blotting and immunohistochemistry were performed to detect the levels of ADAMTS4 and miR-384 in tissues. Luciferase reporter assay, Western blotting and in vitro assay were used to validate that ADAMTS4 was the target gene of miR-384. Neuronal cell line, Neuro-2a, was selected for transfection assay. ADAMTS4 was significantly down-regulated in hippocampi of Alzheimer’s disease mouse model, and negatively correlated with miR-384. Then, ADAMTS4 was identified as a direct target of miR-384. Over-expressing of miR-384 in Neuro-2a showed that ADAMTS4 and the cleaved Reelin fragments were down regulated, and proliferation of neuronal cell lines (Neuro-2a and SH-SY5Y) were inhibited through DAB-1 pathway. In conclusion, these results revealed that miR-384 may play a regulatory role in Reelin via inhibiting ADAMTS4 in neuronal cell lines.

## Introduction

Reelin, a large extracellular glycoprotein, plays important roles both during brain development and in mature brain [1]. Full-length Reelin is a 3460-amino acid precursor protein which consists of eight unique repeats, and its molecular weight is 410 kDa [1,2]. The precursor protein is proteolytically cleaved by metalloproteases such as ADAMTS4 (aggrecanase-1), a member of the A disintegrin and metalloproteinase with thrombospondin motifs (ADAMTSs) family [3]. ADAMTS4 cleaves Reelin at two sites. One occurs at the N-terminal (N-t cleavage) between the second and the third repeat, and the other occurs at the C-terminal (C-t cleavage) between the sixth and seventh repeat. The biological significance of Reelin cleavage is still controversial [4]. Some researchers suggested that Reelin cleavage releases mature fragment from the precursor protein [5], while others supported that the cleavage decreases Reelin activity [6].

An increasing number of studies have showed that altered Reelin expression and processing may contribute to the pathogenesis of several neurodegenerative disorders [7–11], such as Alzheimer’s disease (AD). Previous work by Chin *et al*. [12] found that Reelin-expressing pyramidal cells in the entorhinal cortex significantly decreased in human amyloid precursor protein (hAPP) transgenic mice and humans with AD. Kocherhans *et al.* [13] revealed that reduced Reelin-mediated signaling enhanced amyloid-β plaque formation in aged Reelin-deficient transgenic AD mice. However, the data of Reelin expression in the context of AD is inconsistent [14], and the molecular mechanism of its roles in AD is largely unknown.

MicroRNAs (miRNAs) are endogenous short (~22 nt) RNAs that function as post-translational regulators by binding to the 3’-untranslated region (3’-UTR) of target mRNAs for degradation or translational repression [15]. Among these miRNAs, miR-384 is involved in the metabolic regulation of nerve cells in central nervous system (CNS) [16,17], and has been reported to be altered in several CNS related diseases, such as glioma [18], Parkinson’s Disease (PD) [19] and AD [20].

In this study, by using AD mouse model and two neuronal cell lines, we revealed the regulatory role of miR-384 to ADAMTS4, an important Reelin cleaving enzyme.

## Materials and methods

### Animals

Ten 8-weeks C57BL/6J mice were randomly divided into control group and AD group. Each mouse was placed in a stereotaxic frame after scalp sterilization, skull exposure and incision, and then injected with 1μL of liquid (10g/L Aβ1-42, sigma for AD group, or saline for control group), with bregma coordinates: −1.0 mm lateral, −0.3 mm posterior and −2.5 mm below. After six weeks, mice of both groups were sacrificed and the hippocampi were separated for further studies.

All animal experiments were approved by Animal Ethics Committee of Hubei University of Traditional Chinese Medicine. All procedures involving animals were in compliance with the care and use guidelines of experimental animals established by the Ministry of Agriculture of China.

### Cell culture, miRNA transfection and cell proliferation assay

Cells were cultured in DMEM (HEK293, Neuro-2a) or RPMI 1640 (SH-SY5Y) with 10% FBS (Gibco). Cells grew on petri dishes to a density of 1×10^5^/cm^2^, and then incubated with miR-384 mimics (auuccuagaaauuguucaua, RiboBio, China) w/o negative control mimics (RiboBio, China), by using lipofectamine 2000 (Invitrogen) following the instruction. After 24 hours, cells were digested, diluted in medium and counted. Then, cells were seeded on a 96-well culture plate with concentration of 1×10^4^ cells/well. To estimate the cell proliferation speed, cells of three wells for each transfection condition were digested and counted every day.

### RNA extraction and RT-PCR assay

For RNA extraction, 40 mg tissue sample or 1×10^5^ cultured cells were homogenized in 2 mL Trizol (Invitrogen), and then 1 mL chloroform was added. After vortex and centrifugation at 10,000×g for 10 minutes, the supernatant was collected. Next, double volume of ice-cold RNase-free isopropanol was added to the supernatant, and centrifugated at 12,000×g for 15 minutes. After centrifugation, RNA precipitate was collected and dried at 60°C for 10 minutes. RNA was reversely transcribed into cDNA using Hifair™ II 1st Strand cDNA Synthesis kits (Yeasen, Shanghai, China). RT-PCR were performed by using Hieff™ qPCR kits (Yeasen, Shanghai, China) on CFX96 (Biorad) according to the manufacturer’s instruction. Relative expression levels of genes versus internal reference were calculated by CFX Manager™ software (Bio-Rad). Primers for amplification of ADAMTS4 (m5’-ATGGCCTCAATCCATCCCAG, m5’-AAGCAGGGTTGGAATCTTTGC, h 5’-GAGGAGGAGATCGTGTTTCCA, h5’-CCAGCTCTAGTAGCAGCGTC), b-actin (5’-CCAAGGCCAACCGCGAGAAGATGAC, 5’-AGGGTACATGGTGGTGCCGCCAGAC), miR-384 (5’-CGGCGGTCATTGGTAGAAATTG, 5’-CCAGTGCAGGGTCCGAGGTAT) and U6 (5’-CGGCGGTCGTGAAGCGTTCCAT, 5’-CCAGTGCAGGGTCCGAGGTAT), or for reverse transcription of mRNA (d18T), miR-384 (GTCGTATCCAGTGCAGGGTCCGAGGTATTCGCACTGGATACGACTATGTT) and U6(5’-GTCGTATCCAGTGCAGGGTCCGAGGTATTCGCACTGGATACGACAAAAAT) were synthesized by Tsingke (Wuhan, China).

### Protein extraction and western blot

For protein extraction, 10 mg tissue sample was homogenized in 2 mL RIPA buffer (50mM tris-Cl, pH 8.0, 1% NP-40, 1% Na-deoxycholate, 150mM NaCl, 0.1% SDS, 0.05mM PMSF), or 1×10^5^ cells suspended in 2 mL RIPA buffer. The supernatant was collected by centrifugation at 10,000×g for 10minutes. Protein quantification was tested by bicinchoninic acid protein quantification kit (Yeasen, Shanghai, China). Then the proteins were adjusted to approximate 5 mg/mL using loading buffer, separated by SDS-PAGE and transferred on to 0.22μm PVDF membranes (Millipore). After blocking with 5% skimmed milk, the membranes were incubated at room temperature for 4 h with anti-Reelin (20689-1-AP, Proteintech, 1:1000), anti-ADAMTS4 (SRP00350, Saierbio, 1:800), anti-p-DAB1(P03459, Boster, 1:800), anti-DAB1 (PAB18464, Abnova 1:800) and anti-pAkt (66444-1-Ig, Proteintech 1:800), respectively. Then, the membranes were washed repeatedly (more than five times) using 0.1% TBS-tween-20 buffer, and incubated with 5% skimmed milk containing peroxidase-conjugated goat anti-rabbit and goat anti-mouse antibodies (1:20000) at room temperature for 2 h. Target bands were detected by ECL illumination kit (Millipore, Shanghai, China).

### Plasmid Construction and site-directed mutation

Partial 3’-UTR region of human and mouse ADAMTS4 gene were amplified with primers (5’-CCCCTCGAGATTTAGCACCAGGGAAGGGGA, 5’-AAAGCGGCCGCCACTCATATTTGTCACCTTCT) using cDNA of HEK293 or mouse brain tissue, respectively. Amplified DNAs were cloned into pSiCHECK-2 (Promega, Wuhan, China) by utilizing XhoI and NotI restriction enzyme sites to construct pSi2-hUTR and pSi2-mUTR. To generate mutant plasmids pSi2-hm1 and pSi2-hm2, plasmid pSi2-hUTR was further amplified by primers (5’-AATCCAGGGTGTTGGTGACA, 5’-AGGTAGCTAGGTTAGGGCTATAATGGGTGAG or 5’-TGGAAGCTAGGTTAGGGCTATAATGGGTGAG), phosphorylated by PNK (Fermentas, Wuhan, China), and self linked. All the recombinant plasmids were tested by Sanger sequencing (Tsingke, Wuhan, China).

### Dual-fluorescence assay

Plasmids pSi2-hUTR, pSi2-mUTR, pSipSi2-hm1 and pSi2-hm2 were transfected into HEK293T with 4μM miR-384mimics (RiboBio, China) or 4μM control mimics (RiboBio, China). After culture of more than 48 h, dual-fluorescence assay was performed by Dual-Glo™ Luciferase Assay System (Promega), according to the manufacturer’s instruction.

### Immunohistochemistry (IHC)

The hippocampus tissues were fixed by 4% paraformaldehyde for 7 d, embedded in paraffin, and sectioned to 2-5 μm sections on glass slides. The sections were incubated with anti-ADAMTS4 (SRP00350, Saierbio, Shanghai, China) for 48 h firstly and then with HRP-conjugated goat-anti-rabbit antibody (Proteintech, Shanghai, China). Next, the sections were stained by DAB staining kit (DA1010, Solarbio, Beijing, China) following the instruction, and photographed under microscopy.

### Statistical analysis

All significance tests were carried out by R3.6.1, using unpaired t-test with Welch’s correction. Statistically significant data were indicated by asterisks: * p<0.05, ** p<0.01 or *** p<0.001.

## Results

### miR-384 and ADAMTS4 are negatively correlated in Normal/AD mouse brain

The expression of ADAMTS4 in different tissues was detected by RT-PCR analysis, and the result showed that ADAMTS4 was highly expressed in liver and brain (Fig. 1A). Comparing between the Aβ induced AD mice group and the normal group, it was indicated that expression of ADAMTS4 was inhibited on both mRNA and protein levels in AD mice brains (Fig. 1C and D). Likewise, the result of IHC showed that ADAMTS4 was down-regulated in hippocampi of AD mice (Fig. 1E).

**Figure 1.**
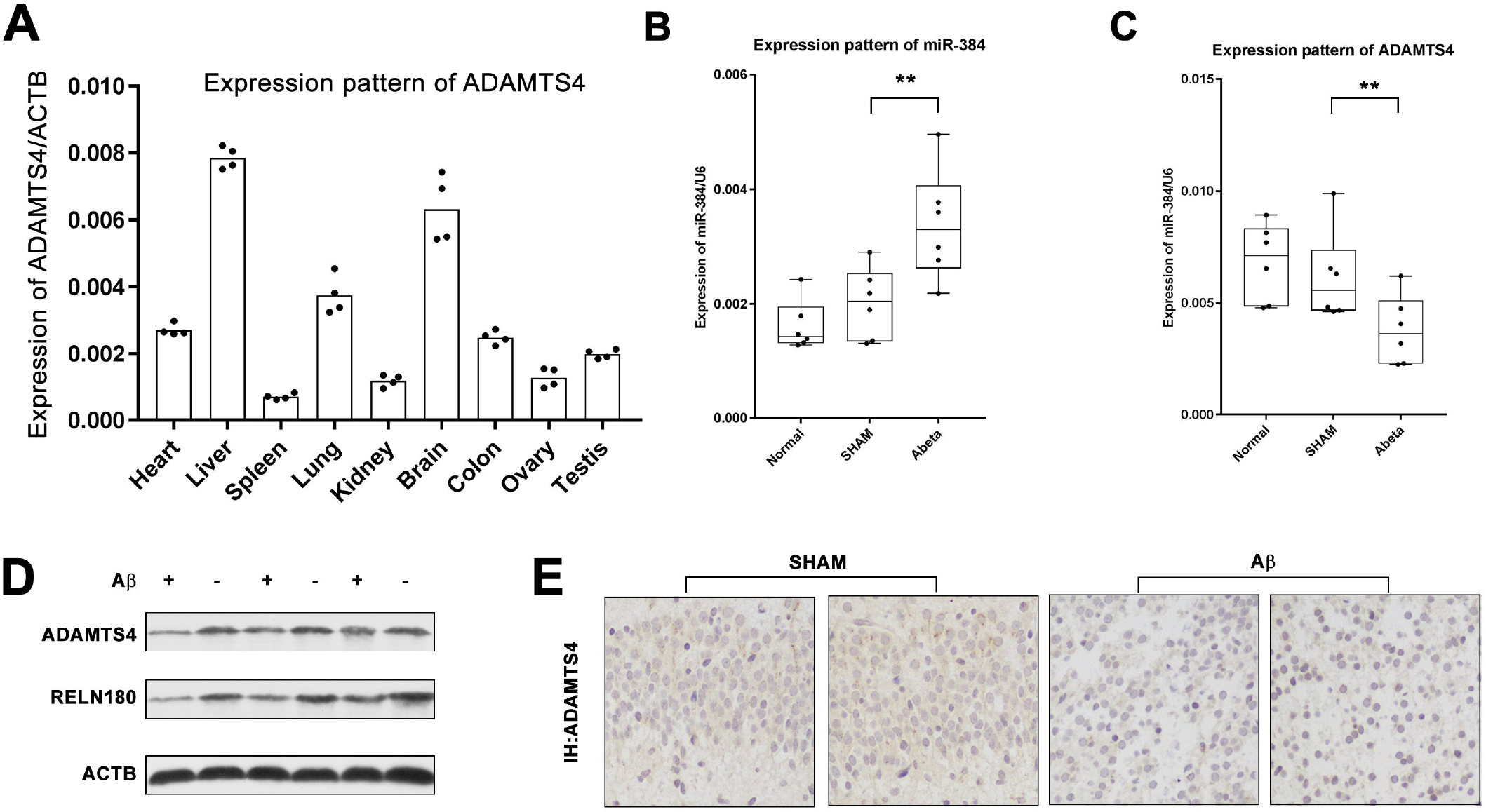
MiR-384 and ADAMTS4 are negatively correlated. (A) Expression level of ADAMTS4 in different mouse tissues was detected by RT-PCR. ADAMTS4 was highly expressed in liver and brain. (B-D) Injection of Aβ_1-42_ in mice hippocampi increased expression level of miR-384 (B), and decreased expression of ADAMTS4 both at mRNA level (C) and protein level (D). Also, the cleaved Reelin fragment (180kD) was down-regulated (D). (E) IHC results of mice hippocampi showed the down-regulation of ADAMTS4 in CA3 region of hippocompi.

Reelin proteolytic cleavage generates Reelin fragments of different molecular weights, with the 180 kDa N-terminal fragment as the predominant. Thus, the cleaved 180 kD Reelin fragment was detected. As compared with the control group, the cleaved 180 kD Reelin fragment was inhibited in AD group (Fig. 1D). The expression of miR-384 was detected, and it was up-regulated AD mice brains (Fig. 1B), which was reversely correlated to ADAMTS4. These results suggested a probability of the interaction between miR-384 and ADAMTS4.

### miR-384 regulates ADAMTS4 by single targeting site

The conserved miR-384 targeting sites of ADAMTS4 (Fig. 2A) were predicted using TargetScan [21]. Plasmids pSi2-hUTR and pSi2-mUTR were constructed, containing human and mouse target sites, respectively. To verify the predicted sites, mutant plasmids pSi2-hm1 and pSi2-hm2 were also constructed. Comparison of Gaussia luciferase and firefly luciferase activities of transfected HEK293 cells showed that miR-384 inhibited Gaussia luciferase through human and mouse 3’-UTR of ADAMTS4, and didn’t inhibit through mutated targeting sites (Fig. 2B). MiR-384 mimics with different concentration were transfected into HEK293, and levels of miR-384 increased correspondingly (Fig. 2C). Meanwhile, the mRNA level of ADAMTS4 was significantly decreased (Fig. 2D). The result of Western blot showed that the protein level of ADAMTS4 was down-regulated by the transfection of miR-384 at a dosage dependent manner. Besides, the cleaved 180 kD Reelin fragment was decreased after miR-384 transfection (Fig. 2E). These results indicated that miR-384 regulates ADAMTS4 through the predicted targeting site.

**Figure 2.**
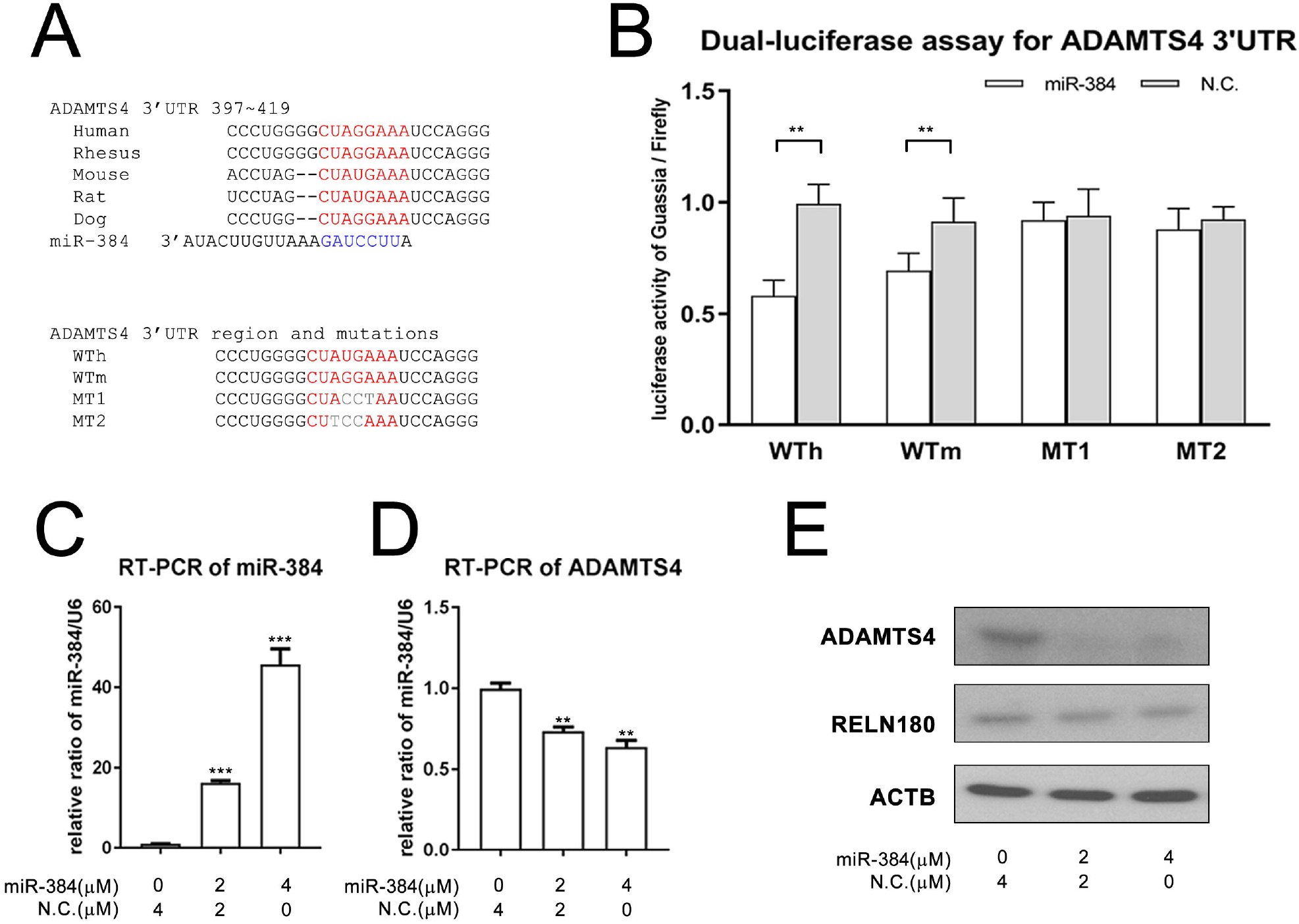
MiR-384 regulates ADAMTS4 by targeting its 3’-UTR. (A) Conserved miR-384 targeting site of 3’-UTR of mammalian ADAMTS4 were predicted by TargetScan, and the mutant sequences were used in this study. (B) Dual-luciferase assays showed that miR-384 transfection resulted in lower Gaussian luciferase activity with plasmids carrying wild type human and mouse target sites, and no changes with plasmids carrying mutated target sites. These results indicated that miR-384 regulates ADAMTS4 through the targeting site of 3’-UTR. (C-D) RT-PCRs showed that transfection of miR-384 mimics resulted in high level of miR-384 in HEK293 cells, and lower mRNA level of ADAMTS4. (E) Western blot results showed that protein levels of ADAMTS4 and the cleaved Reelin (180kD) were both down-regulated.

### miR-384 inhibits Reelin signaling and neuronal cell proliferation

In canonical Reelin signaling pathway, Reelin exerts its functions by bingding to two receptors, apolipoprotein E receptor 2 (ApoER2) and very low-density lipoprotein receptor (VLDLR), and inducing phosphorylation of the intracellular protein Dab1 which activates multiple signal transduction pathways, such as Akt, PI3K and CrK/CrkL. Thus, the concentration of Dab1 and Akt can be used as an indicator of Reelin signaling.

To evaluate the possible functions of miR-384 on Reelin signaling, Neuro-2a cells were transfected with miR-384 mimics, and the cells with over-expressed miR-384 were obtained (Fig. 3A). RT-PCR and Western blot analyses showed that the mRNA and protein levels of ADAMTS4 in Neuro-2a were significantly down-regulated by over-expressing miR-384 (Fig. 3B and E).

**Figure 3.**
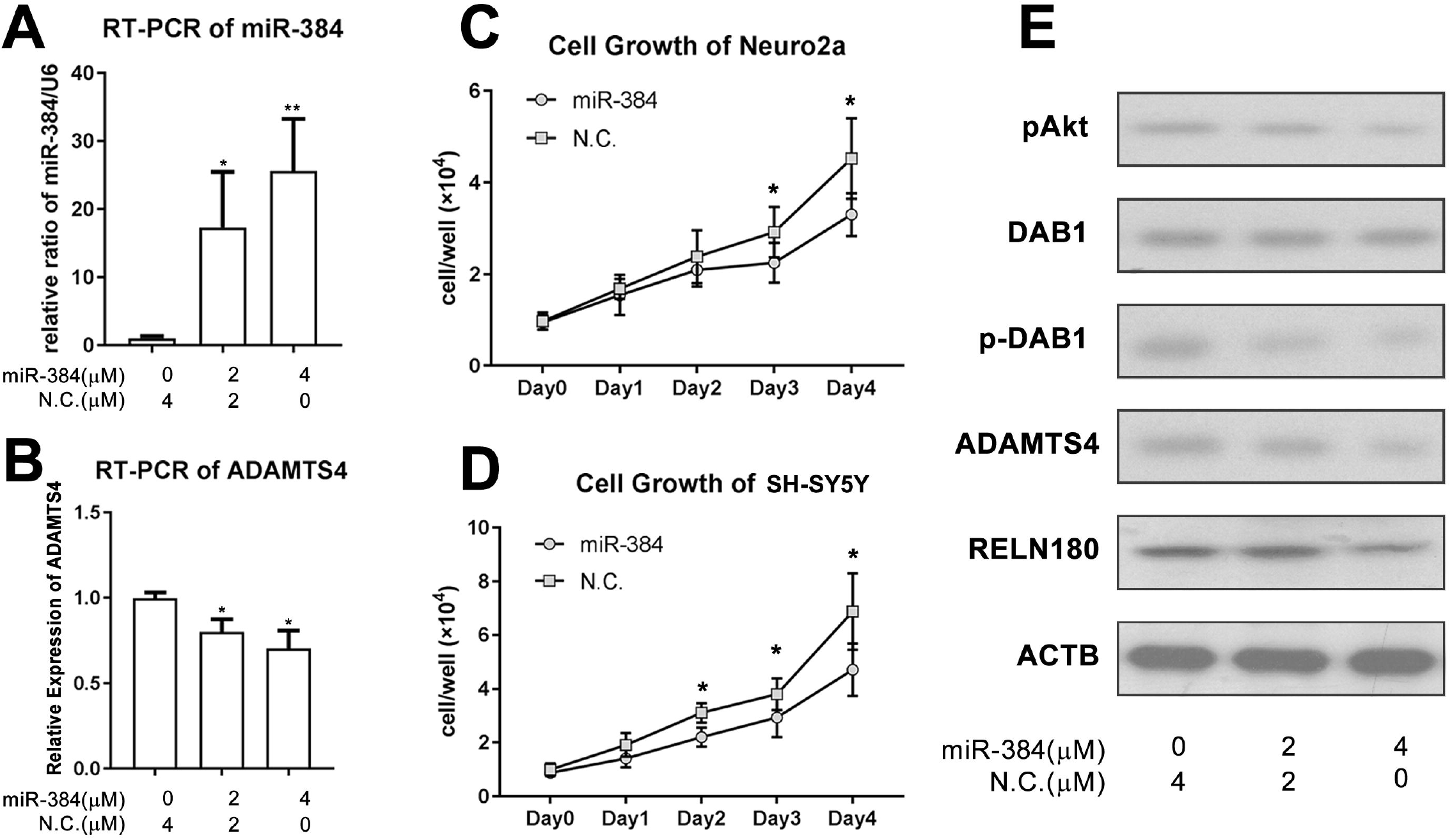
MiR-384 inhibits cell proliferation of neuronal cells. (A) RT-PCRs showed that transfection of miR-384 mimics resulted in high level of miR-384 in Neuro-2a and (B) low mRNA level of ADAMTS4. (C-D) The proliferation ability was investigated by cell growth curve of Neuro-2a and SH-SY5Y. The results showed that over-expressing miR-384 in the cells revealed a significant inhibition in cell proliferation. (E) Results of Western blot analyses showed that protein levels of ADAMTS4, 180kD Reelin fragment, phosphorylated Dab-1 and phosphorylated Akt were down-regulated. These results indicated that miR-384 down-regulated ADAMTS4 and Reelin, and inhibited downstream phosphorylation of Dab-1 and Akt.

Moreover, the 180kD Reelin fragments were significantly decreased, and phosphorylation Dab-1 and Akt were up-regulated, without alteration in total Dab-1 and Akt (Fig. 3E). Also, proliferation assays of Neuro-2a and SH-SY5Y were performed to determine the role of miR-384 in neuronal cell proliferation. Over-expressing miR-384 in the cells revealed a significant inhibition in cell proliferation (Fig. 3C and D). These results suggested that miR-384 may inhibit Reelin signaling and neuronal cell proliferation.

## Discussion

ADAMTS4, a member of ADAMTSs family, has been shown to cleave Reelin, a large secreted glycoprotein which is considered to be involved in the pathogenesis of AD [9–12,14]. Using TargetScan, miR-384 was predicted to be a potential regulator of ADAMTS4. In this study, the relationship between miR-384 and ADAMTS4 was investigated, and the effect they may have on Reelin signaling and neuronal cell proliferation were tested. First, the expression levels of ADAMTS4 and miR-384 in AD mouse group and control group were measured, and the results showed that miR-384 and ADAMTS4 were negatively correlated in the groups. Then, the targeting site of ADAMTS4 by miR-384 was validated using dual-luciferase reporter assay system. Furthermore, the role of miR-384 on Reelin signaling and neuronal cell proliferation was explored, and it was found that miR-384 inhibited Reelin signaling and neuronal cell proliferation.

As a key regulator of gene expression, miRNA is attracting widespread interest in various fields [21], especially in multiple cancer types where it serves as a oncogenic role or a tumor suppressor, depending on its target gene. For miR-384, previous studies also focused on its tumor suppressive role in cancers, such as breast cancer [22], osteosarcoma [23] and glioma [18]. However, it is also involved in the metabolic regulation of nerve cells in CNS and dysregulated in several CNS related diseases. Recently, Liu *et al*. [16] reported that miR-384 regulated chronic cerebral ischemia-induced neuronal apoptosis via a Snhg8/miR-384/Hoxa13/FAM3A axis. The present findings, for the first time, demonstrated that ADAMTS-4 was regulated by miR-384 via direct binding the predicted targeting site, resulting in suppression of ADAMTS-4 at both mRNA and protein level. Furthermore, the interactions among miR-384, ADAMTS4 and Reelin signaling in neuronal cells were explored, and their possible mechanism was demonstrated.

The importance of Reelin signaling pathway in synaptic plasticity and learning and memory has been well-documented [1]. Dysregulation of Reelin is involved in a number of neurodegenerative disorders characterized by cognitive deficits, especially in AD where it may contribute to its pathogenesis [13,20,24]. Therefore, more work is needed to achieve a better understanding of Reelin and its signaling mechanisms, which may throw a light on clinical therapies against cognitive deficits in AD.

## Conclusion

In conclusion, the findings of the present study showed that miR-384 may play a regulatory role in ADAMTS4-Reelin pathway by targeting a single conserved site of ADAMTS4, and over-expression of miR-384 inhibits Reelin signaling.

## Acknowledgements

This work was supported by National Natural Science Foundation of China (Grant Number: 81703954). The authors thank reviewers for their detailed review of this article and the many constructive comments that greatly improved the manuscript.

